# Evolution of a Functionally Intact but Antigenically Distinct DENV Fusion Loop

**DOI:** 10.1101/2023.03.22.533803

**Authors:** Rita M. Meganck, Deanna Zhu, Stephanie Dong, Lisa J. Snoderly-Foster, Yago R. Dalben, Devina Thiono, Laura J. White, Aravinda M. DeSilva, Ralph S. Baric, Longping V. Tse

## Abstract

A hallmark of Dengue virus (DENV) pathogenesis is the potential for antibody-dependent enhancement, which is associated with deadly DENV secondary infection, complicates the identification of correlates of protection, and negatively impacts the safety and efficacy of DENV vaccines. ADE is linked to antibodies targeting the fusion loop (FL) motif of the envelope protein, which is completely conserved in mosquito-borne flaviviruses and required for viral entry and fusion. In the current study, we utilized saturation mutagenesis and directed evolution to engineer a functional variant with a mutated FL (D2-FL) which is not neutralized by FL-targeting monoclonal antibodies. The FL mutations were combined with our previously evolved prM cleavage site to create a mature version of D2-FL (D2-FLM), which evades both prM- and FL-Abs but retains sensitivity to other type-specific and quaternary cross-reactive (CR) Abs. CR serum from heterotypic (DENV4) infected non-human primates (NHP) showed lower neutralization titers against D2-FL and D2-FLM than isogenic wildtype DENV2 while similar neutralization titers were observed in serum from homotypic (DENV2) infected NHP. We propose D2-FL and D2-FLM as valuable tools to delineate CR Ab subtypes in serum as well as an exciting platform for safer live attenuated DENV vaccines suitable for naïve individuals and children.

## INTRODUCTION

Dengue virus (DENV) is a member of the *Flavivirus* genus and is a major global public health threat, with four major serotypes of DENV found worldwide. Dengue causes ∼400 million infections each year, of which ∼20% of cases present clinically, a subset of which may progress to severe Dengue Hemorrhagic Fever/Dengue Shock Syndrome (DHF/DSS).^1, 2^ DENV is transmitted through *Aedes* mosquito vectors, and globalization and global warming are increasing the endemic range of Dengue worldwide.^3, 4^ The pathogenesis of Dengue is complex, as first-time infections are rarely severe and lead to serotype-specific immunity. However, re-infection with a different serotype increases the risk of developing DHF/DSS.^5^ This is thought to be due to the phenomenon of antibody-dependent enhancement (ADE), in which poorly neutralizing cross-reactive (CR) antibodies (Abs) lead to enhanced viral uptake and infection of unique cell populations in an Fcγ-receptor-mediated manner.^6^

ADE remains a major challenge for DENV vaccine development.^7^ The leading DENV vaccine platforms in clinical testing are tetravalent live attenuated virus mixtures of all four serotypes. However, creating formulations that elicit a balanced response has proven challenging.^8^ Additionally, lab-grown strains differ from patient-derived DENVs in both maturation status and antigenicity.^9^ In particular, Abs targeting the fusion loop (FL) have been reported to neutralize lab and patient strains with differing strengths and have been observed to facilitate Fcγ-receptor uptake *in vitro* and therefore ADE.^10–12^ Currently, there is a single FDA-approved DENV vaccine, Dengvaxia. However, it is only approved for use in individuals aged 9-16 with previous DENV infection living in endemic areas and is contraindicated for use in naïve individuals and younger children. In naïve children, vaccination stimulated non-neutralizing CR Abs that increased the risk of severe disease after DENV infection.^13, 14^ Other DENV vaccines have been tested or are currently undergoing clinical trial, but thus far none have been approved for use in the United States.^15^ The vaccine Qdenga has been approved in the European Union, Indonesia, and Brazil, although vaccine efficacy in adults, naïve individuals, and with all serotypes has not yet been shown.

The DENV FL is located in Envelope (E) protein domain II (EDII) and is involved in monomer-monomer contacts with EDIII.^16^ During the DENV infection cycle, low pH triggers a conformational change in the E protein.^17^ The structure of the virion rearranges, and individual monomers form a trimer with all three FLs in the same orientation, ready to initiate membrane fusion.^16, 17^ The core FL motif (DRGWGNGCGLFGK, AA 98-110) is highly conserved, with 100% amino acid conservation in all DENV serotypes and other mosquito-borne flaviviruses, including Yellow fever virus (YFV), Zika virus (ZIKV), West Nile virus (WNV), Kunjin virus (KUNV), Murray Valley encephalitis virus (MVEV), Japanese encephalitis virus (JEV), Usutu virus (USUV), and Saint Louis encephalitis virus (SLEV; Figure 1A). Although the extreme conservation and critical role in entry have led to it being considered extremely difficult to change the FL, we successfully tested the hypothesis that massively parallel directed-evolution could produce viable DENV FL mutants that were still capable of fusion and entry, while altering the antigenic footprint. The FL mutations, in combination with optimized prM cleavage site mutations, ablate neutralization by the prM- and FL-Abs, retain sensitivity to other protective Abs, and provide a novel vaccine strategy for DENV.

**Figure 1:**
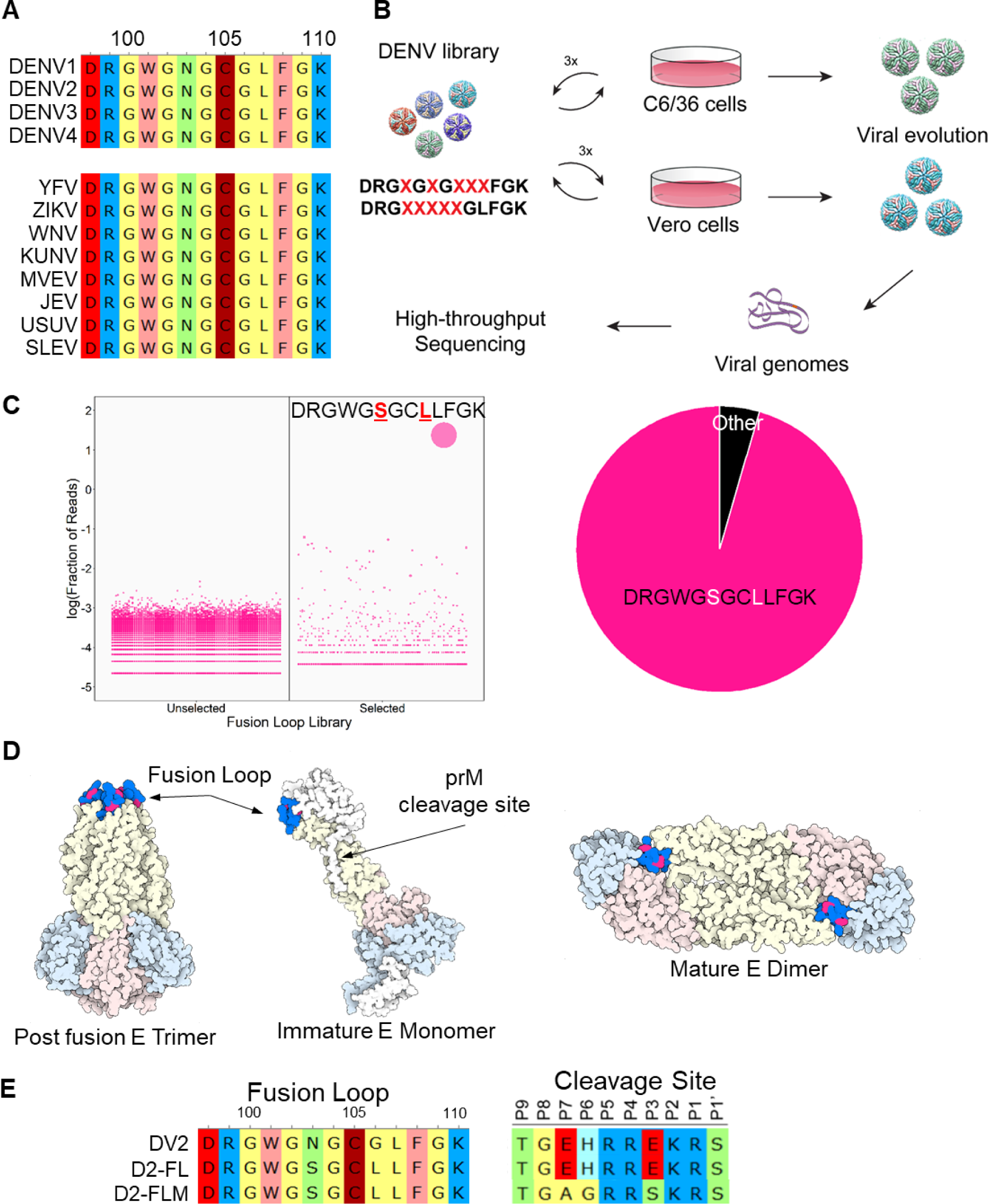
Generation of DENV2 fusion loop mutants via directed evolution. A) Alignment of Top: Dengue virus fusion loops; Bottom: Mosquito-borne flavivirus fusion loops, including Yellow Fever virus (YFV), Zika virus (ZIKV), West Nile virus (WNV), Kunjin virus (KUNV), Murray Valley Encephalitis virus (MVEV), Japanese Encephalitis virus (JEV), Usutu virus (USUV), and Saint Louis Encephalitis virus (SLEV). Amino acids are colored by functional groups: negatively charged (red), positively charged (blue), nonpolar (yellow), polar (green), aromatic (pink), and sulfide (dark red). B) Schematic of directed evolution procedure. Saturation mutagenesis libraries were used to produce viral libraries, which were passaged three times in either C6/36 or Vero 81 cells. At the end of the selection, viral genomes were isolated and mutations were identified by high-throughput sequencing. C) Left: Bubble plot of the sequences identified from either the unselected or selected (passage 3) C6/36 DENV libraries. Right: Pie chart of the sequences from passage 3 C6/36 DENV libraries. D) Structure of the DENV envelope with the fusion loop mutations highlighted in red. E) Sequences of the fusion loop and furin cleavage site of DENV2, D2-FL, and D2-FLM.

## RESULTS

To engineer a virus with a novel antigenic footprint at the FL, we targeted the core conserved FL motif. We generated two different saturation mutagenesis libraries, each with 5 randomized amino acids: DRG**X**G**X**G**XXX**FGK (Library 1; AA 101, 103, 105-107) and DRG**XXXXX**GLFGK (Library 2 AA 101-105). Library 1 was designed to mutate known residues targeted by FL mAbs while Library 2 focused on a continuous linear peptide that is the epitope for FL-Abs to maximally alter antigenicity.^18^ Saturation mutagenesis plasmid libraries were used to produce viral libraries in either C6/36 (*Aedes albopictus* mosquito) or Vero 81 (African green monkey) cells and passaged three times in their respective cell types. Following directed evolution, viral genomes were extracted and subjected to deep sequencing to identify surviving and enriched variants (Figure 1B). Due to the high level of conservation, it was not surprising that most mutational combinations failed to yield viable progeny. In fact, evolutions carried out on Library 2 only yielded wild-type sequences. For Library 1, wild-type sequences dominated in Vero 81 evolved libraries. However, a novel variant emerged in C6/36 cells with two amino acid changes: DRGWG**<u>S</u>**GC**<u>L</u>**LFGK. The major variant comprised ∼95% of the population, while the next most populous variant (DRGWG**<u>S</u>**GC**<u>W</u>**LFGK) comprised only 0.25% (Figure 1C). Bulk Sanger sequencing revealed an additional Env-T171A mutation outside of the FL region. Residues W101, C105, and L107 were preserved in our final sequence, supporting the importance of these residues.^16^ When modeled on the pre-fusion DENV2 structure, the N103S and G106L mutations are located at the interface with the neighboring monomer EDIII domain, protected from the aqueous environment. In the post-fusion form, the two residues are located between W101 and F108 and form the bowl concavity above the chlorine ion in the post-fusion trimer (Figure 1D). We used reverse genetics to re-derive the FL N103S/G106L/T171A mutant, which we term D2-FL. As enhancing Abs also target prM,^19^ we also created a mature version of D2-FL termed D2-FLM, containing both the evolved FL motif and our previously published evolved prM furin cleavage site, which results in a more mature virion like those found in infected patients (Figure 1E).^9, 20^

We performed growth curves comparing DENV2, D2-FL, and D2-FLM in both C6/36 and Vero 81 cells. In C6/36 cells, the growth of all three viruses was comparable, reaching high titers of 10^6^-10^7^ FFU/mL. However, in Vero 81 cells, both FL mutant viruses were highly attenuated, with a 2-2.5 log reduction in titer (Figure 2A). The species-specific phenotype in culture involved a change from insect to mammalian cells, as well as a change in growth temperature. To investigate if the mutant viruses were more unstable at higher temperatures, we performed a thermostability assay, comparing viruses incubated at temperatures ranging from 4-55°C before infection. The three viruses had comparable thermostabilities, indicating that this does not explain the attenuation of the FL mutants (Figure 2B). Because the D2-FLM virus contains mutations that increase prM cleavage frequency, we also assayed the maturation status of the three viruses by western blot. D2-FL had a comparable prM:E ratio to the isogenic wildtype DENV2 (DV2-WT), while, as expected, D2-FLM had a reduced prM:E ratio, indicating a higher degree of maturation (Figure 2C).

**Figure 2:**
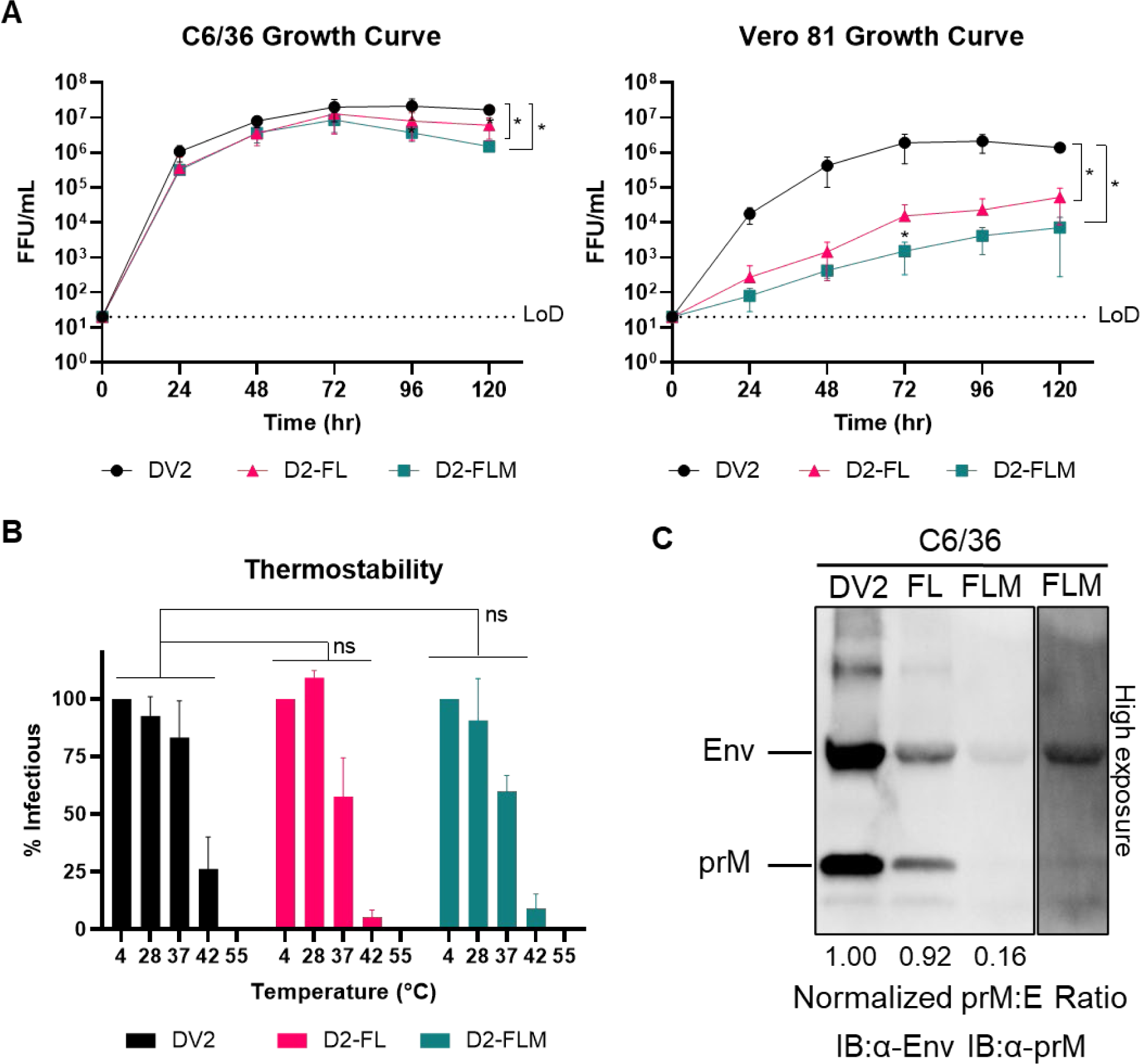
Biological and physical properties of mature DENV2 fusion loop mutants. A) Multistep growth curves (MOI = 0.05-0.1) of DV2-WT, D2-FL, and D2-FLM on C6/36 cells (left) or Vero 81 cells (right). B) Thermostability assay on DV2-WT, D2-FL, and D2-FLM. C) Western blot of virions, blotted against Envelope and prM proteins. The prM:E ratio was determined and normalized to the DV2-WT ratio. Averages of 3 biological replicates are shown. Two-way ANOVA was used for statistical comparison of growth curves and thermostability: ns = not significant; * < 0.05; ** < 0.005; *** < 0.0005.

Next, we characterized the ability of Abs targeting the FL to recognize DV2-WT, D2-FL, and D2-FLM with a panel of monoclonal antibodies (mAbs). Importantly, D2-FL and D2-FLM were resistant to mAbs targeting the FL. Neutralization by 1M7 is reduced by ∼2-logs in both variants, 1N5 neutralization is reduced by ∼1-log for D2-FL and reduced to background levels for D2-FL, and no neutralization was observed for 1L6 or 4G2 for either variant (Figure 3A).^18^ Focusing on the D2-FLM virus containing both evolved motifs, we then characterized the antigenicity of the whole virion with a panel of mAbs. As expected, D2-FLM was unable to be neutralized by the prM Abs 1E16 and 5M22; the Ab 2H2 does not neutralize either DV2-WT or D2-FLM (Figure 3B). For Abs targeting epitopes in non-mutated regions, including the ED1 and EDE epitopes that target EDII and EDIII, FRNT_50_ values were generally comparable, although EDE1-C10 shows a moderate but statistically significant reduction between DV2-WT and D2-FLM, indicating that the overall virion structural integrity was intact (Figure 3B).

**Figure 3:**
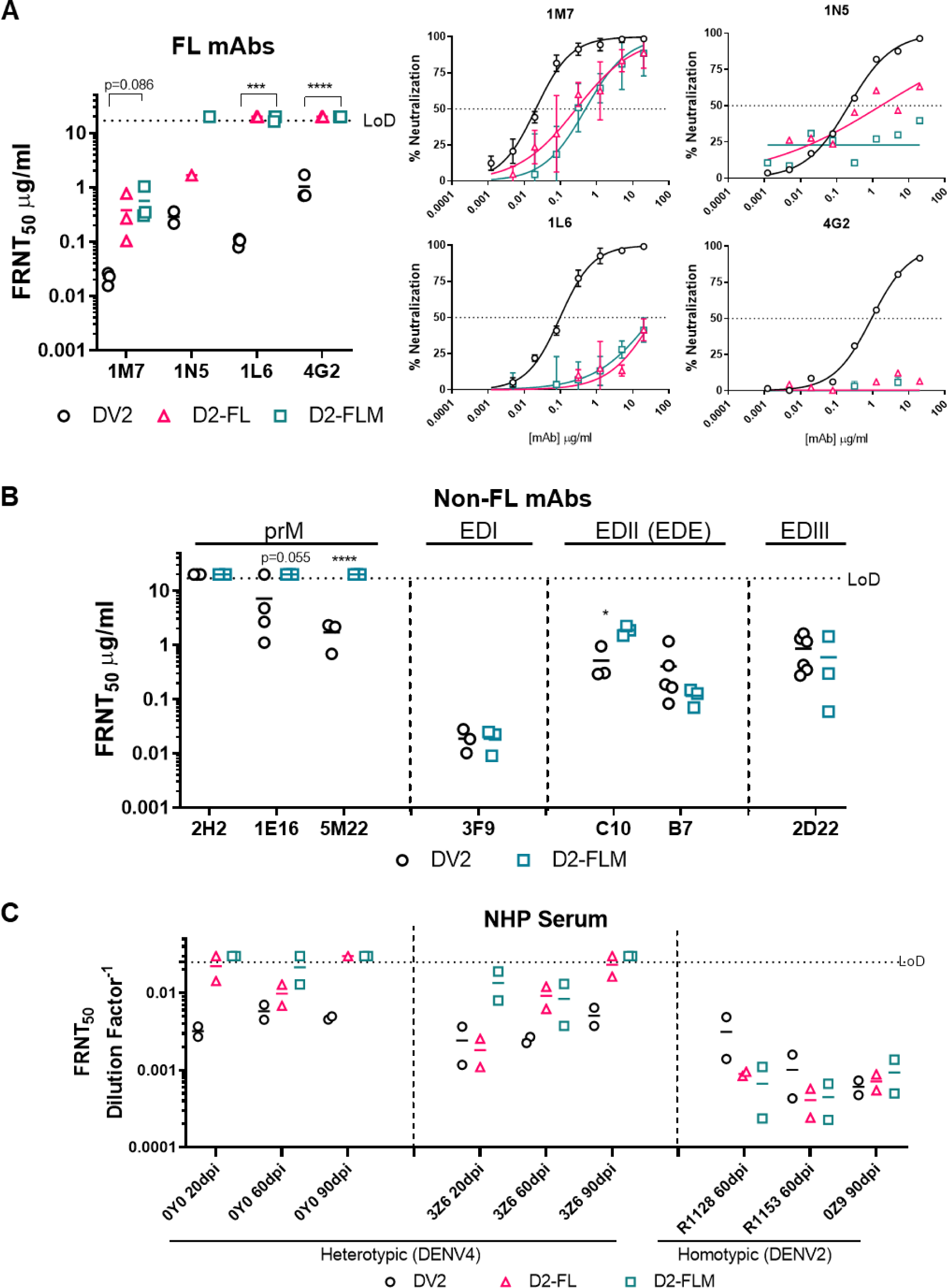
Fusion loop mutant is insensitive to fusion loop mAbs, the major target for cross-reactive Abs in NHPs. A) Left: FRNT_50_ values for neutralization of DV2-WT, D2-FL, and D2-FLM with mAbs against the FL (1M7, 1N5, 1L6, 4G2). All Abs were tested in at least n=3 independent experiments, except 1N5 due to limited Ab. Right: Average neutralization curves for neutralization of DV2-WT, D2-FL, and D2-FLM with mAbs against the DENV2 fusion loop. B) FRNT_50_ values for neutralization of DV2-WT and D2-FLM with mAbs against DENV2 prM (2H2, 1E16, 5M22), EDI (3F9), EDE (C10, B7), and EDIII (2D22). All Abs were tested in at least n=3 independent experiments. C) Neutralization of DV2-WT, D2-FL, and D2-FLM with sera from NHPs infected with either DENV4 or DENV2. FRNT_50_s were compared using Student’s t-test. Significant symbols are as follows: *, P < 0.05; **, P< 0.005; ***, P < 0.0005; ****, P < 0.00005. The data are graphed as means ± standard deviations.

Next, we analyzed neutralization of the D2-FLM virus using serum derived from convalescent humans and experimental infected non-human primates (NHPs). Overall, we tested serum from 6 humans and 9 NHPs at different time points with a total of 27 samples. Serum from a homotypic infected NHP (n=3) did not display a difference in neutralization between DV2-WT, D2-FL, and D2-FLM, confirming that prM and FL epitopes are not significant contributors to the homotypic type-specific (TS) neutralizing Ab response in primates (Figure 3C). In heterotypic vaccination and infection, most of the serum (18/24) did not cross-neutralize (FRNT_50_ < 1:40) DV2-WT, confirming the serotypic difference of DENVs (Table 1). However, in two NHPs infected with DENV4, strong neutralization potency (FRNT_50_ between 1: 100 – 1:1,000) was demonstrated against DV2-WT (Figure 3C). Heterologous cross-neutralization was significantly reduced to background levels (FRNT_50_ < 1:40) against the D2-FLM virus at 90 days post-infection (dpi). Of note, one DENV4 animal (3Z6) showed low levels of neutralization against D2-FLM at early time points (20- and 60-days post-infection), which was eventually lost at later time points. Neutralization observed against D2-FL, in general, fell between DV2-WT and D2-FLM. Interestingly, in animal 3Z6, at 20 dpi, neutralization against DENV-FL was comparable to DV2-WT, while D2-FLM was greatly reduced, indicating that antibodies in the sera targeting the immature virion formed a large portion of the CR response. In contrast, animal 0Y0 displayed less difference in neutralization between D2-FL and D2-FLM, suggesting that FL antibodies were more prominent in this animal. These data suggest that after a single infection, much of the CR Ab responses target prM and the FL and tend to wane over time (Figure 3C). The collection of FL, mature, and FL-mature variants provides new opportunity to delineate antibody composition in complex polyclonal serum from DENV natural infection and vaccination.

**Table 1:**
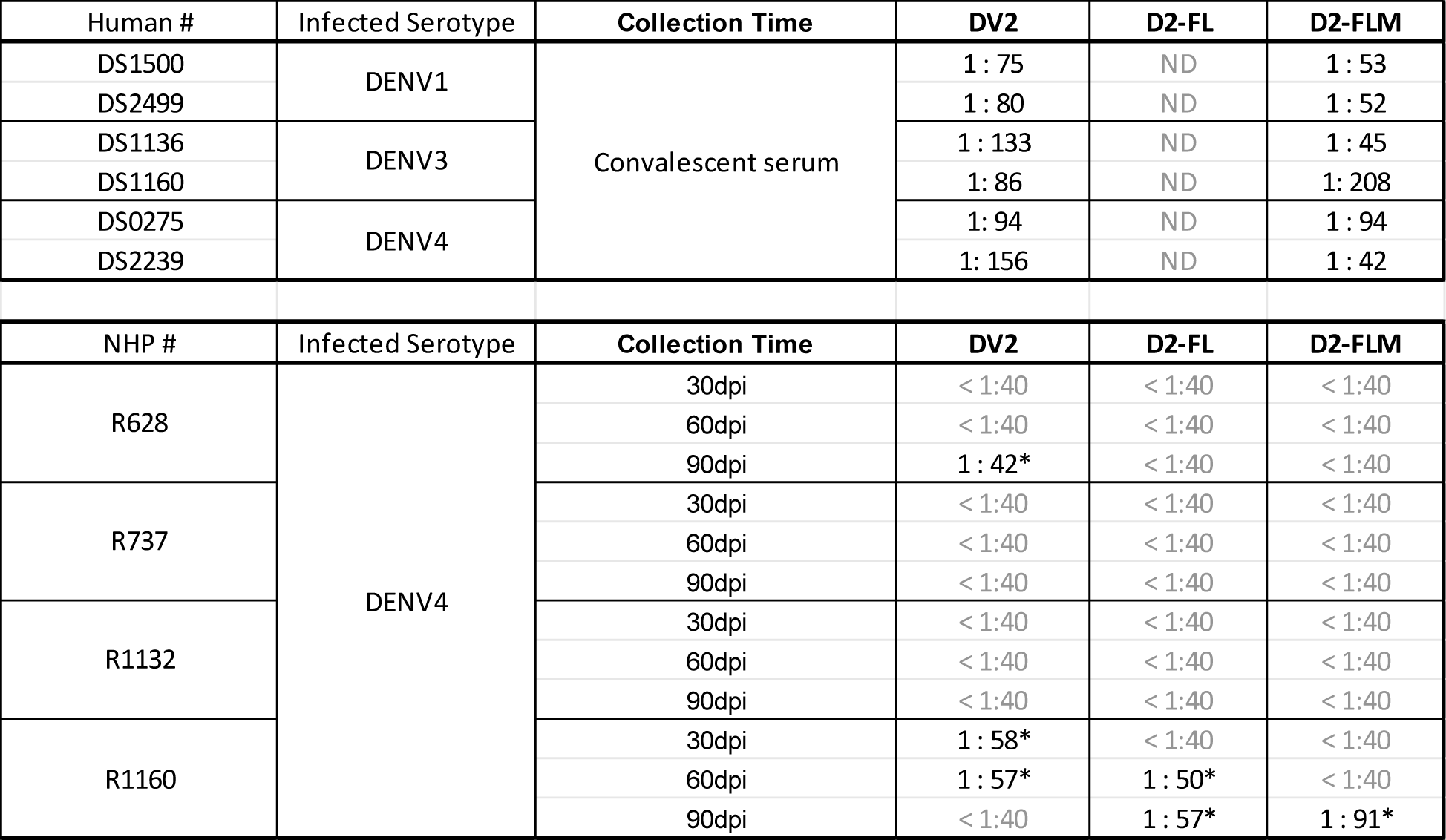
Summary of FRNT_50_s of human convalescent serum and NHP infection serum against DV2, D2-FL and D2-FLM.

## DISCUSSION

Mechanistic understanding of vaccine protection and identification of correlates of protection are immensely important for DENV vaccine development. The dual protective and enhancing properties of DENV Abs create major challenges for dissecting the role of various Ab populations in disease protection. Cross-reactive weakly neutralizing prM- and FL-Abs are often immunodominant after primary DENV infection,^10, 21–24^ and can lead to overestimation of the levels of heterotypic protection in traditional neutralization assays. Since these same antibodies are also associated with ADE,^19^ inaccurate conclusions could have dire consequences if protection *in vitro* translates to the enhancement of disease in human vaccinees. Unfortunately, Ab profiling in polyclonal serum is mainly performed by ELISA, and a neutralization assay that can discriminate Abs does not exist. The D2-FLM variant is not neutralized by FL- and prM-mAbs and appears insensitive to neutralization by these Abs in polyclonal serum. Of note, EDE1-C10 neutralization potency was also reduced in our FL-M variant, further indicating our mutations are affecting the tip of the EDII region which partially overlaps with the EDE1 epitope^25^. In combination with other chimeric DENVs,^26–28^ D2-FLM provides a reagent to distinguish between TS, protective CR (e.g. E dimer epitope, EDE),^29, 30^ and ADE-prone CR (e.g. FL and prM)^24^ Ab subclasses in neutralization assays after infection and vaccination.

Due to the ADE properties of DENV Abs, studies to understand and eliminate ADE phenotype are under active investigation. For example, mAbs can be engineered to eliminate binding to the Fcγ receptor, abolishing ADE.^31^ While this methodology holds potential for Ab therapeutic development and passive immunization strategies, it is not relevant for vaccination. As FL and prM targeting Abs are the major species demonstrated to cause ADE *in vitro* and are thought to be responsible for ADE-driven negative outcomes after primary infection and vaccination,^10–12, 32^ we propose that genetic ablation of the FL and prM epitopes in vaccine strains will minimize the production of these subclasses of Abs responsible for undesirable vaccine responses. Indeed, efforts have been made to reduce the availability of the FL or reduce the ability of FL Abs to drive ADE. Covalently locked E-dimers and E-dimers with FL mutations have been engineered as subunit vaccines that reduce the availability of the FL, thereby reducing the production of FL Abs.^33–36^ DENV subunit vaccines are an area of active study;^37^ however, monomer/dimer subunits can also expose additional, interior-facing epitopes not normally exposed to the cell. Furthermore, dimer subunits are not a complete representation of the DENV virion which presents other structurally important interfaces such as the 3-fold and 5-fold symmetries. Concerns about balanced immunity to all four serotypes also apply to subunit vaccine platforms. Given the complexity of the immune response to DENV, live virus vaccine platforms have thus far been more successful. However, the fusion loop is strongly mutationally intolerant. Previous reported mutations in the fusion loop of mosquito-borne flaviviruses include a L107F mutation in WNV ^38^ and JEV associated with attenuation, and mutations at position 106 in ZIKV(G106A) and DENV(G106V) which were tolerated. Interestingly, we also recovered a mutation in position 106 (G106L) Using directed-evolution, we successfully generated our D2-FLM variant that combines viability with the desired Ab responses. Therefore, the D2-FLM variant is a novel candidate for a vaccine strain which presents all the native structures and complex symmetries of DENV necessary for T-cell mediated responses and which can elicit more optimal protective Ab responses.^39–41^

Other considerations of high importance when designing a live DENV vaccine include strain selection and serotype balance.^42^ In the current study we used DENV2 S16803, a prototype for DENV2.^43^ However, S16803 was isolated several decades ago, and it may be beneficial to utilize more contemporaneous strains.^8, 44^ Work is currently ongoing to demonstrate the portability of the evolved FL motif on additional DENV2 strains and other serotypes, which is essential for tetravalent vaccine production. D2-FLM was highly attenuated in Vero cells, creating a challenge for vaccine production. Therefore, further adaptation of this strain to grow efficiently in mammalian cells while retaining its antigenic properties is needed. Taken together, the FLM variant holds exciting new possibilities for a new generation of DENV vaccines, as well as a platform to readily measure TS and CR ADE-type responses and thereby assess the true protective potential of any DENV vaccine trials and safeguard approval of DENV vaccines for human use.

## ACKNOWLEDGEMENTS

We thank members of the Tse, Baric, and DeSilva laboratories for helpful discussions. This work was supported by NIAID R01AI107731 to A.D. and R.S.B., P01AI106695 to R.S.B., NIAID F30AI160898 to D.R.Z. L.J.W. is supported by NIAID P01 5112869.

## AUTHOR CONTRIBUTION

R.M.M. and L.V.T. designed the study. R.M.M. performed high-throughput sequencing preparation and analysis. R.M.M., D.Z., S.D., L.J.S., Y.D., and L.V.T. performed experiments. D.T., L.J.W., A.M.D.S., and R.S.B. provided reagents. L.V.T. and R.S.B. provided oversight of the project and funding. R.M.M. wrote the manuscript. L.V.T. reviewed and revised the final version. All authors approved the final version of the manuscript.

## MATERIALS AND METHODS

### Cells and viruses

C6/36 (ATCC CRL-1660) were grown in MEM (Gibco) with 5% FBS (HyClone), 1% penicillin/streptomycin (Gibco), 0.1mM nonessential amino acids (Gibco), 1% HEPES (Gibco), and 2mM GlutaMAX (Gibco), cultured at 32°C with 5% CO_2_. Vero 81 cells (ATCC CCL-81) were grown in DMEM/F12 (Gibco) with 10% FBS, 1% penicillin/streptomycin (Gibco), 0.1mM nonessential amino acids (Gibco), and 1% HEPES (Gibco), cultured at 37°C with 5% CO_2_. DENV viruses were grown in C6/36 or Vero 81 cells maintained in infection media. C6/36 infection media consists of Opti-MEM (Gibco) with 2% FBS (HyClone), 1% penicillin/streptomycin (Gibco), 0.1mM nonessential amino acids (Gibco), 1% HEPES (Gibco), and 2mM GlutaMAX (Gibco). Vero 81 infection media consists of DMEM/F12 (Gibco) with 2% FBS, 1% penicillin/streptomycin (Gibco), 0.1mM nonessential amino acids (Gibco), and 1% HEPES (Gibco). DENV2 strain S16803 was used in this study.^43^ Sequences used for the alignments include DENV1 WestPac-74 (U88535.1), DENV2 S-16803 (GU289914.1), DENV3 3001 (JQ411814.1), DENV4 Sri Lanka-92 (KJ160504.1), YFV 17D (NC_002031.1), SLEV Kern217 (NC_007580.2), JEV (NC_001437.1), USUV Vienna-2001 (NC_006551.1), MVEV (NC_000943.1), WNV-1 NY99 (NC_009942.1), and ZIKV MR-766 (NC_012532.1).

### DENV Reverse Genetics

DENV2 S16803 was used in this study. Recombinant viruses were created using a four-plasmid system as previously described,^45^ consisting of the DENV genome split into four segments, each cloned into a separate plasmid. The DENV plasmids were digested and ligated to form a single template for *in vitro* transcription. The resulting RNA was electroporated into either C6/36 or Vero cells. Virus-containing supernatant was harvested at 4-5 days post electroporation and passaged. DENV variants were created through site-directed mutagenesis of the DENV plasmids.

### Library Generation and Directed Evolution

DENV fusion loop libraries were generated through saturation mutagenesis of the indicated resides, based on a previously published protocol.^20, 46^ Degenerate NNK oligonucleotides were used to amplify the region, generating a library of mutated DNA fragments. Q5 DNA Polymerase was used with less than 18 cycles to maintain accuracy. The resulting library was cloned into the DENV reverse genetics system. The ligated plasmids were electroporated into DH10B ElectroMax cells (Invitrogen) and directly plated on 5,245mm^2^ dishes (Corning) to avoid bias from suspension culture. Colonies were pooled and purified using a Maxiprep kit (Qiagen), and the plasmid library used for DENV reverse genetics (above). Viral libraries were passaged three times in the corresponding cell type.

### High-throughput Sequencing and Analysis

Viral RNA was isolated with a QIAamp viral RNA kit (Qiagen), and cDNA produced using the Superscript IV Reverse Transcriptase (Invitrogen). Amplicons were prepared for sequencing using the Illumina TruSeq system with two rounds of PCR using Q5 Hot Start DNA polymerase (NEB). For the first round of PCR, primers were specific to the DENV2 E sequence surrounding the fusion loop motif with overhangs for the Illumina adapters. After purification, this product was used as the template for the second round of PCR using Illumina P5 and P7 primers containing 8-nucleotide indexes. Purified PCR products were analyzed on a Bioanalyzer (Agilent Technologies) and quantified on a Qubit 4 fluorometer (Invitrogen). Amplicon libraries were run on a MiSeq system with 2×150bp reads. Plasmid and P0 libraries were sequenced at a depth of ∼4.5 million reads; later passages were sequenced at a depth of ∼750,000 reads. Custom perl and R scripts were used to analyze and plot the data as previously published.^20^

### DENV Growth Kinetics

One day before infection, 5×10^5^ cells were seeded in every well of a 6-well plate. Cells with infected with an MOI of 0.05-0.1, estimating 1×10^6^ cells on the day of infection. Infection was carried out for one hour in the incubator, followed by 3x washes with PBS and replenishment with fresh infection medium. 300 uL of viral supernatant was collected at 0, 24, 48, 72, 96, and 120 hours and stored at -80°C. All experiments were performed independently at least three times.

### DENV Focus-Forming Assay

Titers of viral supernatant were determined using a standard DENV focus-forming assay. In brief, cells were seeded at 2×10^4^ cells per well of a 96-well plate one day before infection. The next day, 50 uL of 10-fold serial dilution of viral supernatant were added to each well for 1 hour in the incubator. After, 125uL of overlay (Opti-MEM, 2% FBS, NEAA, P/S, and methylcellulose) was added to each well. Infection was allowed to continue for 48 hours in the incubator. Overlay was removed, and each well rinsed 3x with PBS followed by a 30 minute fixation with 10% formalin in PBS. Cells were blocked in permeabilization buffer (eBioscience) with 5% nonfat dried milk. Primary Abs anti-prM 2H2 and anti-E 4G2 from nonpurified hybridoma supernatant were used at a 1:500 dilution in blocking buffer. Goat anti-mouse HRP secondary (SeraCare KPL) were used at a 1:1000 dilution in blocking buffer. Followed washing, foci were developed using TrueBlue HRP substrate (SeraCare) and counted using an automated Immunospot analyzer (Cellular Technology).

### Thermal Stability Assay

The indicated viruses were thawed and incubated at temperatures ranging from 4°C to 55°C for one hour. Following, viral titers were determined by focus-forming assay as described above.

### Western Blotting

Viral supernatants were combined with 4X Laemmli Sample Buffer (Bio-Rad) and boiled at 95°C for 5 minutes. After SDS-PAGE electrophoresis, samples were transferred to PVDF membrane and blocked in 3% nonfat milk in PBS-T. A polyclonal rabbit anti-prM (1:1000; Invitrogen PA5-34966) and polyclonal rabbit anti-Env (Invitrogen PA5-32246) in 2% BSA in PBS-T were incubated on the blot for 1 hour at 37C. Goat anti-rabbit HRP (1:10,000 Jackson ImmunoLab) in 3% milk in PBS-T was incubated on the blot for 1 hour at room temperature. Blots were developed by SuperSignal West Pico Plus chemiluminescent substrate (ThermoFisher). Blots were imaged on an iBright FL1500 imaging system (Invitrogen). The pixel intensity of individual bands was measured using ImageJ, and the relative maturation was calculated by using the following equation: (prMExp/EnvExp)/(prMWT/EnvWT). All experiments were performed independently a minimum of three times.

### FRNT Assay

Focus reduction neuralization titer (FRNT) assays were performed as described previously with C6/36 cells.^20^ 1×10^5^ cells were seeded in a 96-well plate the day prior to infection. Abs or sera were serially diluted and mixed with virus (∼100 FFU/well) at a 1:1 volume and incubated for 1 hour in the incubator. The mixture was added onto the plate with cells and incubated for 1 hour in the incubator, then overlay was added (see Focus-Forming Assay) and plates were incubated for 48 hours. Viral foci were stained and counted as described above (Focus-Forming Assay). A variable slope sigmoidal dose-response curve was fitted to the data, and values were calculated with top or bottom restraints of 100 and 0 using GraphPad Prism version 9.0. All experiments were performed independently at least two times, due to limited amounts of human serum.

### Statistical Analysis

GraphPad Prism version 9.0 was used for statistical analysis. Titer and % infection of D2-FL and D2-FLM were compared to the DV2 using two-way ANOVA. FRNT_50_s were compared using Student’s t-test. Significant symbols are as follows: *, P < 0.05; **, P< 0.005; ***, P < 0.0005; ****, P < 0.00005. The data are graphed as means ± standard deviations.

